# Bioorthogonal pro-metabolites for profiling short chain fatty acylation

**DOI:** 10.1101/097360

**Authors:** Wilson R. Sinclair, Thomas T. Zengeya, Julie M. Garlick, Andrew J. Worth, Ian A. Blair, Nathaniel W. Snyder, Jordan L. Meier

## Abstract

Short chain fatty acids (SCFAs) play a central role in health and disease. One function of these signaling molecules is to serve as precursors for short chain fatty acylation, a class of metabolically-derived posttranslational modifications (PTMs) that are established by lysine acetyltransferases (KATs) and lysine deacetylases (KDACs). Via this mechanism, short chain fatty acylation serves as an integrated reporter of metabolism as well as KAT and KDAC activity, and has the potential to illuminate the role of these processes in disease. However, few methods to study short chain fatty acylation exist. Here we report a bioorthogonal pro-metabolite strategy for profiling short chain fatty acylation in living cells. Inspired by the dietary component tributyrin, we synthesized a panel of ester-caged bioorthogonal short chain fatty acids. Cellular evaluation of these agents led to the discovery of an azido-ester that is metabolized to its cognate acyl-coenzyme A (CoA) and affords robust protein labeling profiles. We comprehensively characterize the metabolic dependence, toxicity, and histone deacetylase (HDAC) inhibitory activity of these bioorthogonal pro-metabolites, and apply an optimized probe to identify novel candidate protein targets of short chain fatty acids in cells. Our studies showcase the utility of bioorthogonal pro-metabolites for unbiased profiling of cellular protein acylation, and suggest new approaches for studying the signaling functions of SCFAs in differentiation and disease.

## Introduction

Short chain fatty acids (SCFAs) play a critical role as mediators of health and disease.^1–3^ For example, butyrate stimulates differentiation in many cellular models of cancer^4–5^ and displays context-dependent effects on tumorigenesis^6–8^ and immune cell function^9–10^ in whole organisms. In addition to their role in acetyl-coenzyme A (CoA) biosynthesis via the β-oxidation pathway and activity as histone deacetylase (HDAC) inhibitors^11–13^, SCFAs also constitute the requisite precursor for protein short chain fatty acylation. This family of posttranslational modifications (PTMs) derives from the metabolism of SCFAs to acyl-CoAs, which may then be used as cofactors by lysine acetyltransferase (KAT) enzymes (Figure 1).^14–16^ Due to the unique reliance of short chain fatty acylation on both metabolism and KAT activity, this PTM has significant potential to provide insights into the role of these pathways in disease. However, few methods for studying short chain fatty acylation exist. Antibody-based analyses of these modifications are limited by the poor affinity and selectivity of antibodies generated against marks such as butyryl- and propionyl-lysine. This may be why these approaches have, to date, identified only a handful of short chain fatty acylated targets.^17–19^ Bioorthogonal metabolic tracing has emerged as an alternative, unbiased strategy for monitoring SCFA-derived acylation.^20–21^ Pioneered by Hang and coworkers, this approach monitors short chain fatty acylation by using alkynyl-SCFAs to introduce a latent affinity handle into SCFA-modified proteins, enabling their facile visualization and enrichment^22^. A challenge of this approach is that bioorthogonal SCFA tracers must compete with endogenous SCFAs for metabolism and KAT utilization, potentially limiting their efficacy. Thus, the development of highly sensitive bioorthogonal reporters for profiling short chain fatty acylation remains an important goal.

**Figure 1.**
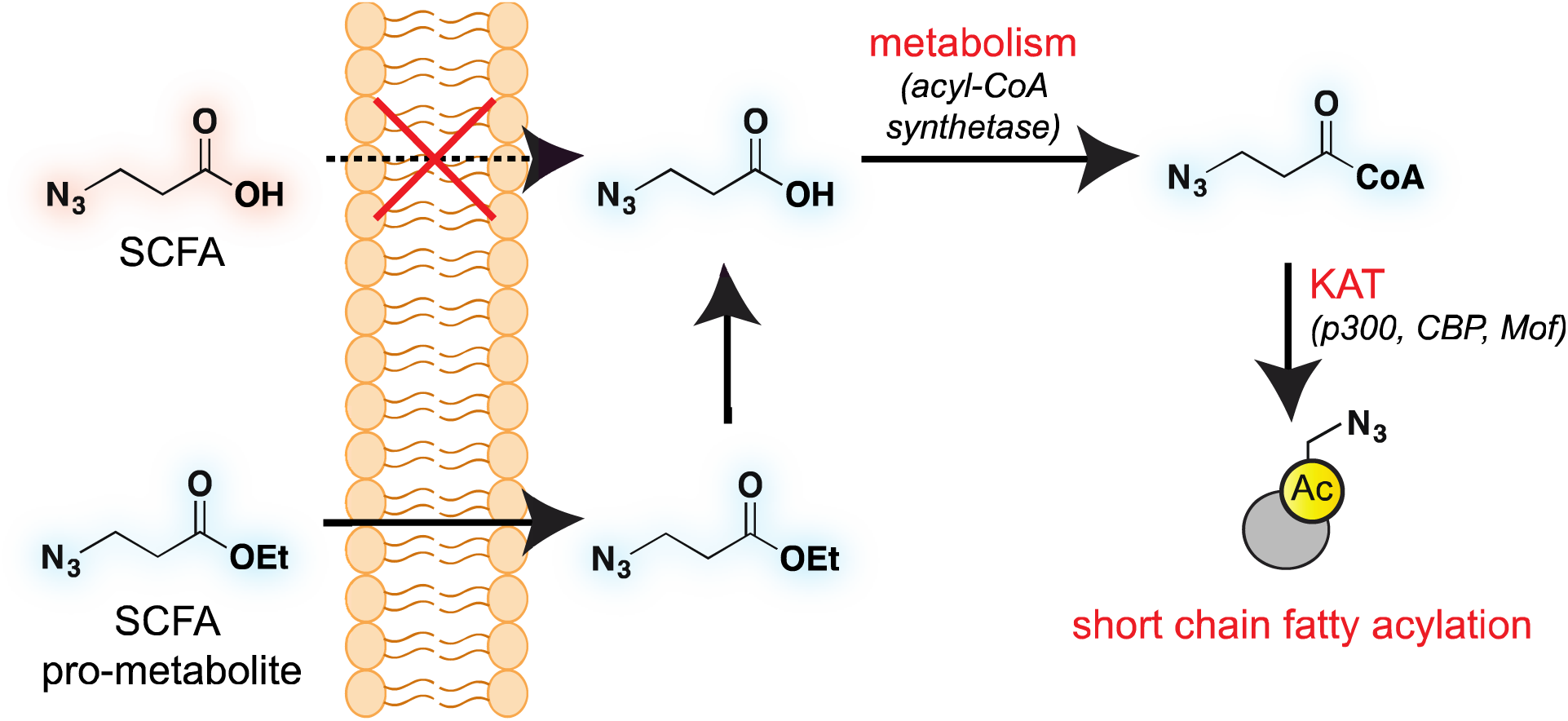
Bioorthogonal pro-metabolite strategy for profiling short chain fatty acylation.

A validated strategy to increase the cellular delivery of small molecules and metabolic tracers is through reversible masking of polar functional groups.^23–25^ Regarding SCFAs specifically, a number of groups have found butyrate’s biological effects to be increased upon masking its polar carboxylate moiety with an ester.^26–27^ In analogy with pro-drugs, we refer to such molecules as “prometabolites.”^28^ A prototypical SCFA pro-metabolite is tributyrin, a naturally-occurring butyrate ester that displays potent antineoplastic and 2Q chemopreventative activity.^29^ Inspired by tributyrin and other pro-metabolites, we hypothesized that masking the polar carboxylate of bioorthogonal SCFAs may similarly increase their potency. Such optimized agents have the potential to provide a robust reporter of short chain fatty acylation, and by proxy, metabolism and cellular KAT/KDAC activity. Towards the goal of defining acylation-dependent signaling in cancer, here we report a bioorthogonal pro-metabolite strategy for profiling short chain fatty acylation (Figure 1). First, we systematically define the influence of SCFA-ester linkage and bioorthogonal chemotype on cellular protein labeling. Next, we demonstrate the ability of optimized agents to form bioorthogonal acyl-CoAs in cells and establish their cell-type and metabolism-specific labeling properties. Finally, we demonstrate the utility of bioorthogonal pro-metabolites to identify novel targets of SCFAs in cells. These studies highlight the power of pro-metabolite approaches to expand the scope of bioorthogonal metabolic tracing methods, and suggest novel strategies for studying the signaling role of short chain fatty acids in health and disease.

## Results and discussion

### Design of a bioorthogonal pro-metabolite library

To identify sensitive agents for profiling short chain fatty acylation, we focused on exploring three structural elements of bioorthogonal SCFAs (Figure 2). First we varied the bioorthogonal reporter, reasoning azides (**1-8**) and alkynes (**9-16**) may demonstrate differential metabolic processing and detection sensitivities. Second, we varied the length of the alkyl chain between the SCFA carboxylate and bioorthogonal reporter. Previous studies have found SCFA chain length to be a crucial factor in the formation of SCFA-CoAs by acyl-CoA synthetases^30–32^, and the utilization of these SCFA-CoAs by KATs.^15, 33^ Third, we varied the carboxylate ester moiety. Here we focused on simple ethyl esters (**2, 6, 10, 14**), as well as carnitine and triacylglycerol esters (Figure 2). Carnitine esters (**3, 7, 11, 15**) were designed to facilitate delivery of SCFAs to the mitochondria, an organelle that shows robust acyl-CoA metabolism.^34^ In contrast, tributyrin-inspired triacylglycerol esters (**4, 8, 12, 16**) were designed to take advantage of the known increased cellular uptake of highly lipophilic molecules, as well as the ability of these esters to be specifically cleaved by endogenous triacylglycerol lipases. An additional advantage of these molecules relative to other esters is their ability to enable suprastoichiometric delivery (three equivalents) of their cognate bioorthogonal SCFAs. This panel of pro-metabolites was readily synthesized using straightforward ester bond formation from the parent bioorthogonal carboxylates (Scheme S1). While the majority of bioorthogonal SCFA derivatives were obtained in good yield, we found carnitine esters to be the most difficult targets due to their challenging purification. Similar optimization was required to obtain triacylglycerol SCFAs, as literature conditions yielded a complex mixture of mono-, di-, and triacylglycerides. These studies establish a systematic panel of bioorthogonal SCFAs to define the structure-activity relationship of pro-metabolites in living cells.

**Figure 2.**
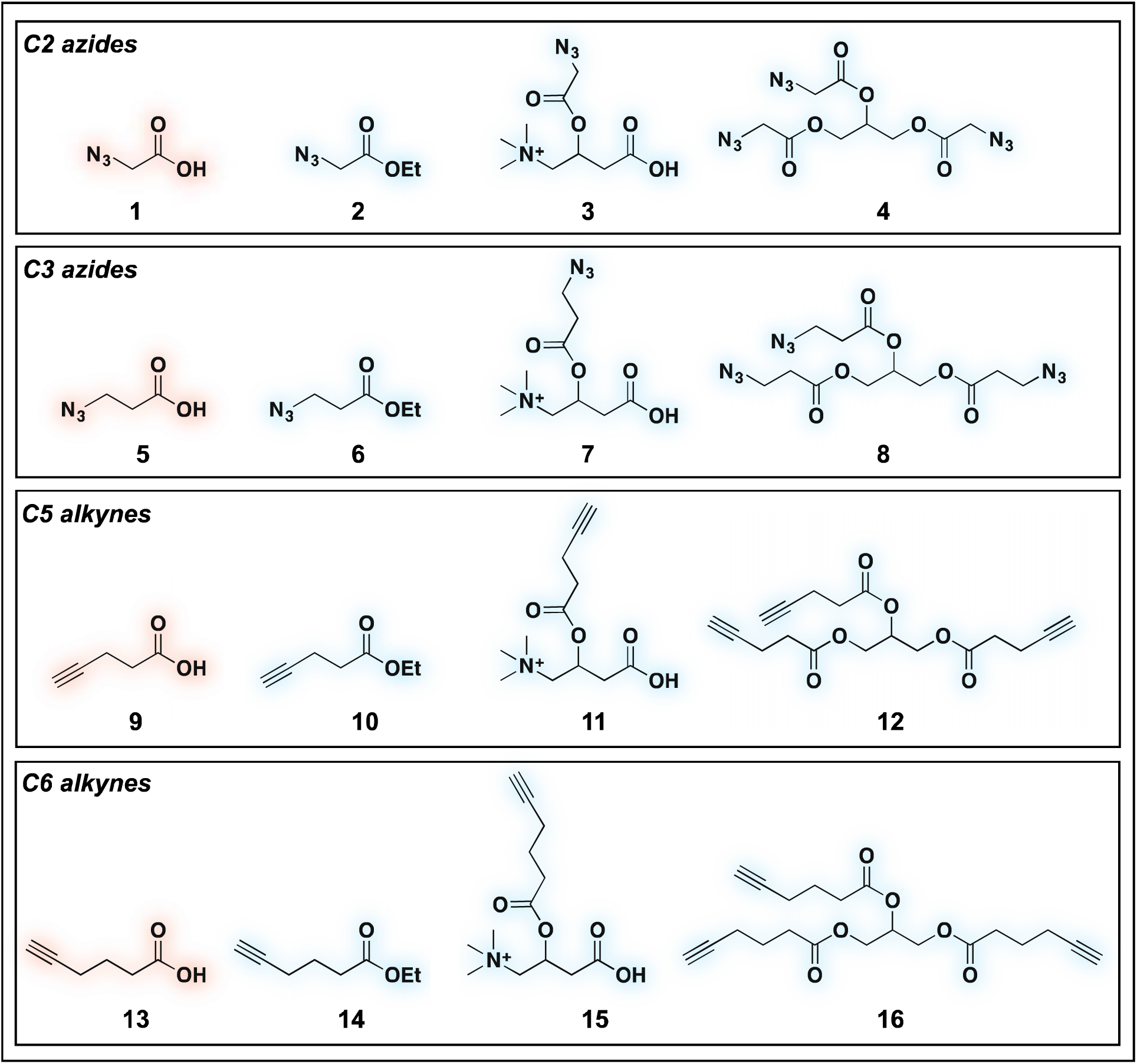
Panel of SCFA pro-metabolites synthesized and evaluated in this study.

### Evaluation of bioorthogonal pro-metabolites for in vitro protein labeling

With these compounds in hand, we next evaluated our library for protein labeling in HepG2 human hepatocellular carcinoma cells (Figure 3). Our choice of liver cancer as a model for evaluation of our pro-metabolites stems from the fact that the liver is a major site of SCFA metabolism in vivo, where it utilizes abundant acyl-CoA synthetase activities to transform SCFAs into acyl-CoAs that support short chain fatty acylation.^35^ Accordingly, we treated HepG2 cells with each of our pro-metabolites at a single concentration for 24 hours. Cells lysates were harvested, subjected to click chemistry with a fluorescent azide or alkyne, and analyzed by SDS-PAGE (Figure 3a). Consistent with previous reports, detection of azide-labeled proteins using a fluorophore alkyne manifested uniformly higher background labeling.^36^ Pro-metabolites demonstrating the most intense protein labeling profiles were the bioorthogonal triacylglycerides (**4, 8, 12, 16**) as well as ethyl azidopropionate (**6**) (Figure 3b-c). Labeling by triacylglycerides **8** and **12** was accompanied by noticeable toxicity at 24 hours (Figure S1a). Interestingly, analysis of the bioorthogonal alkyne series revealed pentynoate esters to be consistently more toxic than their hexynoate counterparts (Figure S1b). This did not correlate with protein labeling or HDAC inhibitory activity, (Figure 3, S2), suggesting pentynoate may manifest its toxic affects through alternative mechanisms. Of note, 5-pentynoic acid structurally resembles 5-pentenoic acid, a cytotoxic agent known to form electrophilic species and inhibit fatty acid oxidation in cells.^37^ Analysis of the active azide pro-metabolites (**4, 6, 8**) caused us to focus on **6** as a lead pro-metabolite. Key rationale for this choice included the fact that **6** provides the most intense labeling of any bioorthogonal SCFA precursor examined (Figure 2b), and robustly labels proteins at concentrations that do not impede cell growth (Figure S1c-d). Acid extraction and LC-MS metabolomics analysis of cells treated with 6 confirmed formation of azidopropionyl-CoA, consistent with the ability of 6 to form cofactors for protein short chain fatty acylation reactions (Figure S3). To our knowledge this represents first direct evidence of bioorthogonal acyl-CoA formation in living cells. These data, combined its facile synthesis, prompted us to further evaluate **6** as an optimized reporter of cellular short chain fatty acylation.

**Figure 3.**
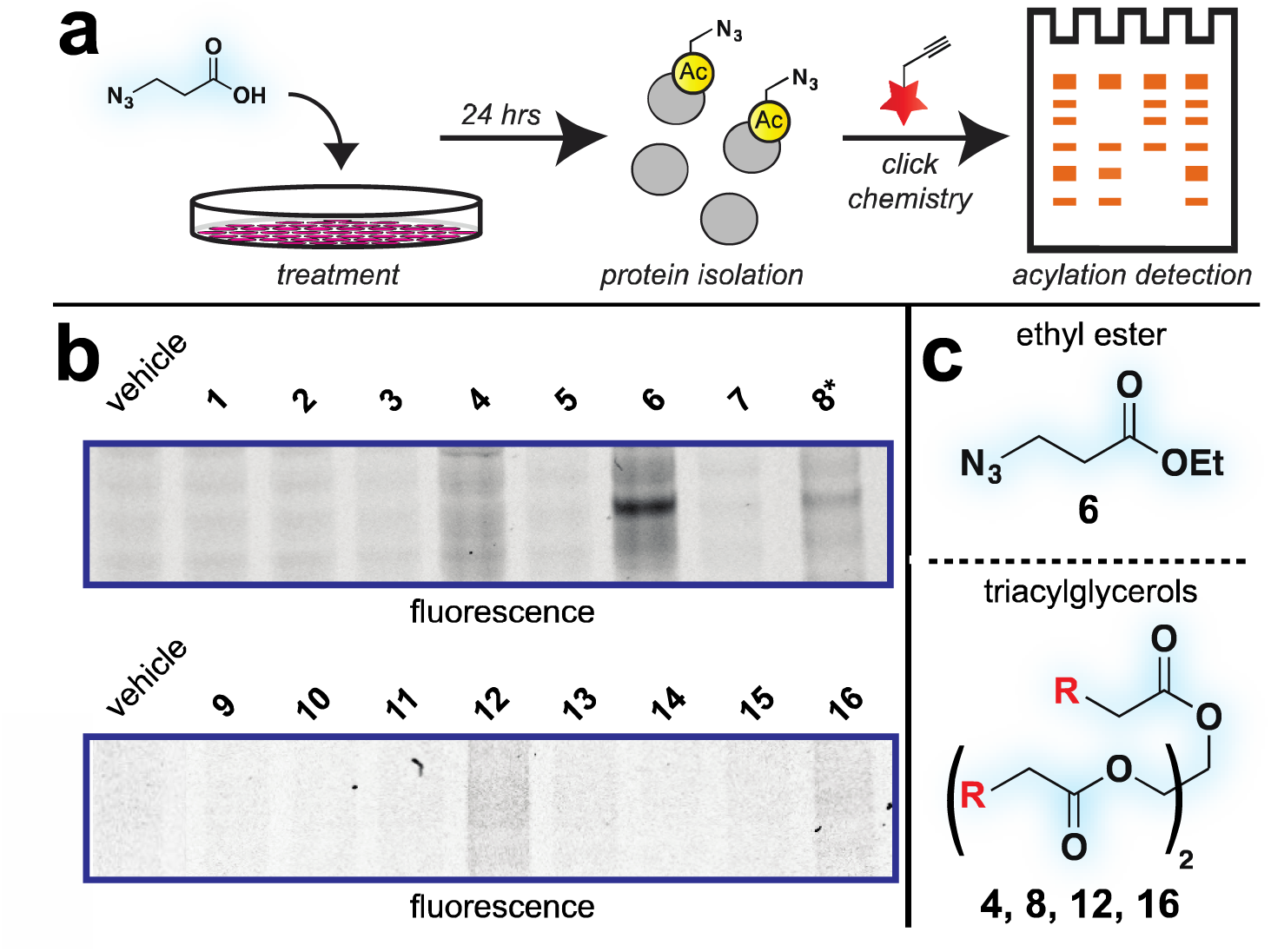
Cellular evaluation of bioorthogonal SCFA pro-metabolites. a) Scheme for cellular analysis and assay of pro-metabolite incorporation via fluorescent gel-based assay. b) Protein labeling by pro-metabolites and parent carboxylates (2.5 mM, 24 h). Asterisk indicates pro-metabolites for which toxicity was observed under these conditions. c) Most active bioorthogonal SCFA pro-metabolite scaffolds.

### Mechanistic analysis of a lead pro-metabolite for short chain fatty acylation profiling

To better define the activity of pro-metabolite **6**, we next evaluated it in a series of experiments using our fluorescent gel-based assay. Cellular labeling of proteins by **6** was dose-dependent, with significant labeling observed at concentrations as low as 0.5 mM (Figure 4a). Analyzing the time-dependent activity of **6** found labeling to peak between 6 and 9 hours (Figure S1d). Of note, the reduced labeling observed at 24 hours suggests short chain fatty acylation implemented by **6** may be reversible, consistent with the known ability of many HDAC enzymes to deacetylate fatty acyl-lysine derivatives.^38–39^ To assess the generality of pro-metabolite **6**, we compsared cell labeling in HepG2 to labeling in cell lines derived from a variety of tissues of origin, including lung (A549) and kidney (HEK-293). Interestingly, each cell line demonstrated intense, but unique, patterns of protein labeling by **6** (Figure 4b). This suggests the ability of **6** to function as a useful reporter of short chain fatty acylation in a number of tissues and cell types.

**Figure 4.**
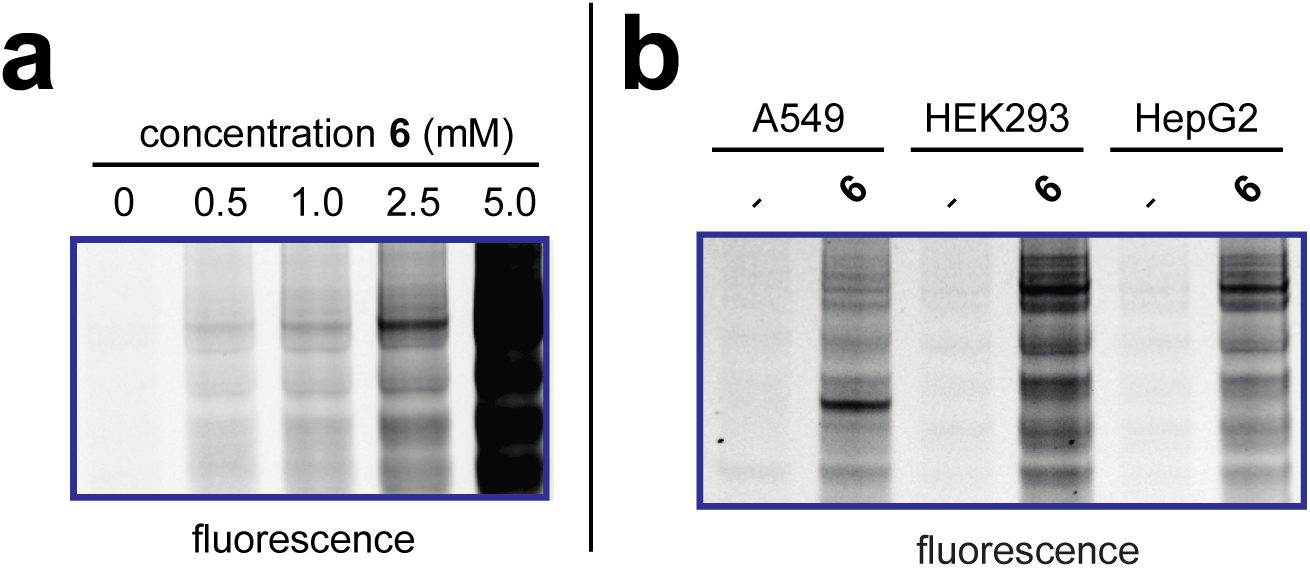
Cellular evaluation of lead pro-metabolite **6**. (a) Dose-dependent labeling of HEK-293T cells. Cells were treated with **6** for 6 h. (b) Cell-line specific protein labeling by pro-metabolite **6**.

Next, we sought to better understand the metabolic determinants of short chain fatty acylation by **6** (Figure 5). First, we explored whether labeling of proteins by **6** was sensitive to competition by endogenous short chain fatty acids such as acetate. Our hypothesis was that acetate may compete with **6** for acyl-CoA synthetase active sites,^35^ thereby reducing bioorthogonal acyl-CoA biosynthesis and competing for sites of protein labeling (Figure 5a). Consistent with this hypothesis, we observed decreased labeling of cellular proteins by **6** upon co-administration of acetate (Figure 5b). Next, we tested the effect of media glucose on labeling on short chain fatty acylation by **6**. Previous studies have found that depriving cells of glucose causes increased utilization of SCFAs for energy production via fatty acid oxidation.^6^ Importantly, this bioenergetic shift requires increased biosynthesis of SCFA-CoAs, and should therefore increase labeling by **6** (Figure 5a) Indeed, we observed increased labeling of cellular proteins by **6** when cells were grown in low glucose compared to nutrient replete media (Figure 5c). Finally, we explored the impact of acetylation dynamics on metabolic labeling. Due to the lack of well-validated chemical probes of KAT enzymes,^40–41^ we focused on assessing how labeling of proteins by **6** was impacted by HDAC inhibitors. Treatment of cells with the HDAC inhibitor SAHA was found to upregulate global acetylation, while coordinately decreasing labeling of cellular proteins by **6** (Figure 5d). This indicates active deacetylation may be required to liberate lysines for short chain fatty acylation by bioorthogonal acyl-CoAs. These studies define pro-metabolite **6** as a general reporter of SCFA metabolism and acylation.

**Figure 5.**
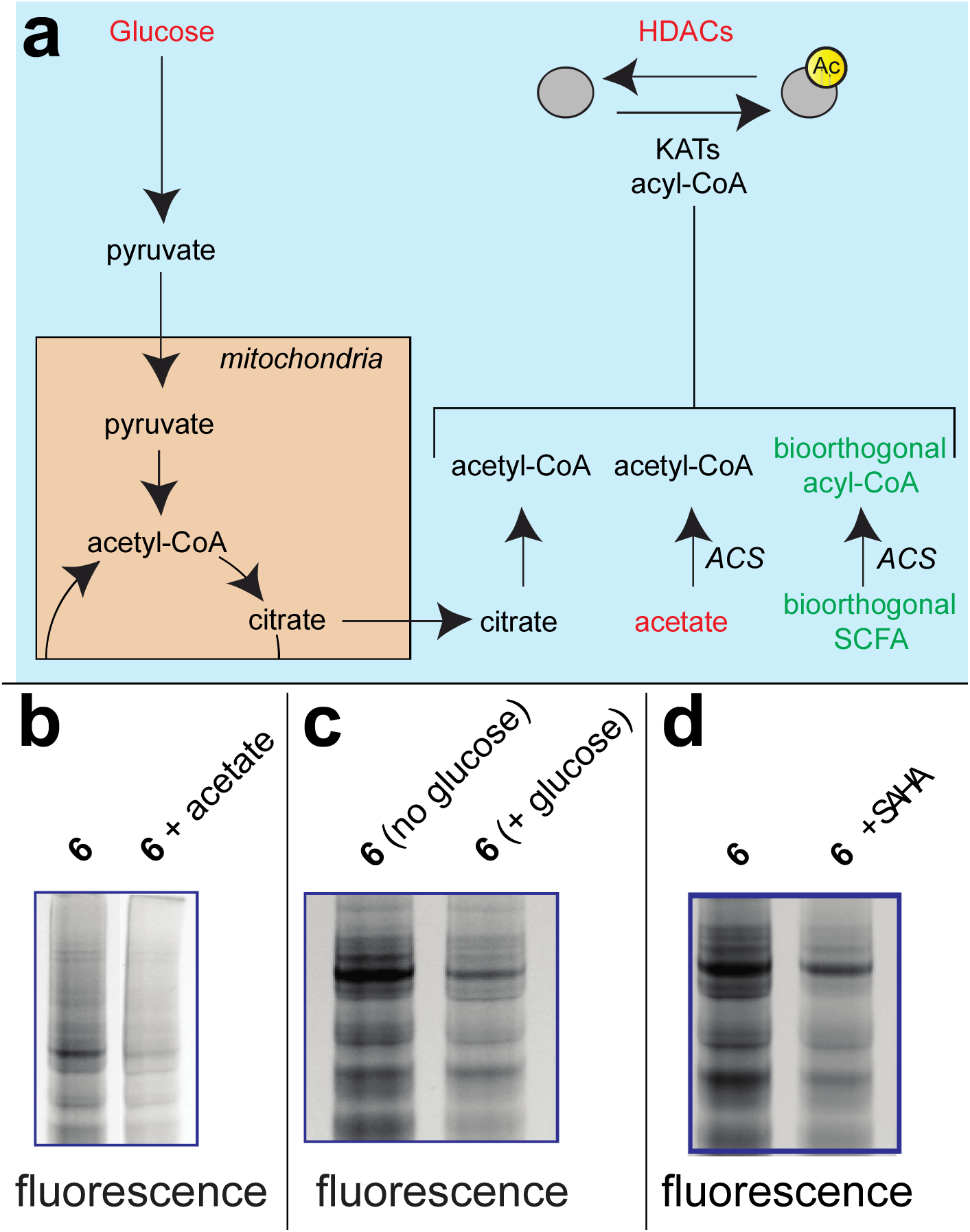
Pro-metabolites are metabolism and HDAC-dependent labeling agents. (a) Schematic depicting biosynthetic origins of acetyl-CoA and SCFA-CoAs used in lysine acylation reactions. Agents in red are manipulated in these studies. ACS = acyl-CoA synthetase enzymes. (b) Protein labeling by bioorthogonal reporter **6** is competed by exogenous SCFAs. (c) Protein labeling by bioorthogonal reporter **6** is enhanced in the absence of glucose, which favors ACS activity. (d) Protein labeling by bioorthogonal reporter **6** is decreased by pre-treatment of cells with the HDAC inhibitor SAHA.

### Profiling the cellular targets of bioorthogonal SCFAs

Finally, we sought to demonstrate the ability of **6** to identify specific targets of short chain fatty acylation in living cells. As a proof-of-concept, we exposed human embryonic kidney cells to either a pulse of **6** or vehicle DMSO for six hours. Following isolation of cell lysates, proteins labeled by **6** were chemoselectively ligated to a biotin alkyne via Cu-catalyzed [3+2] cycloaddition, and enriched over streptavidin-agarose. Enriched proteins were subjected to an on-bead tryptic digest and identified by LC-MS/MS. Candidate short chain fatty acylation targets were defined as proteins enriched greater than 2-fold from cells treated with **6**, relative to a vehicle control (Table S1). Analyzing the abundance of these targets via spectral counting identified 447 low abundance proteins (1-5 spectral counts), 121 medium abundance proteins (5-10 spectral counts), and 91 highly abundant proteins (>10 spectral counts). Of note, many of these proteins, including 41% of highly abundant enriched proteins, have previously been identified as targets of short chain fatty acylation by Hang and coworkers (Table S1).^22^ Gene ontology analysis also supported the utility of **6** as a bioorthogonal reporter, with acetylation being the most strongly enriched term (P = 1.5 × 10^−153^) (Table S2). This analysis also revealed targets of **6** were enriched in nuclear proteins, consistent with the nuclear localization of many KATs (Table S3). Of note, KATs identified in triplicate datasets include EP300, which has previously been observed to catalyze short chain fatty acylation. EP300 may be a target of this modification via autoacetylation.^17, 42–45^ We defer a more detailed validation and discussion of these candidate SCFA targets for future studies. However, overall these findings support the ability of **6** to be applied in global proteomic studies of short chain fatty acylation.

## Conclusions

Here we have described a bioorthogonal pro-metabolite strategy for profiling cellular short chain fatty acylation. Specifically, we find that several alkyne- and azide- SCFAs function as effective protein labeling reagents when delivered as triglyceride analogues (**4, 8, 12, 16**), or as ethyl esters (**6**). Detailed characterization of lead pro-metabolite **6** found that it can form bioorthogonal acyl-CoAs in cells, is active in several cell models, and labels proteins in a manner reflective of cellular metabolism and acetylation dynamics. Our studies suggest that pro-metabolite strategies, which have previously been used to optimize incorporation of azide- and alkyne-containing glycans,^24–25^ may have broad utility in the delivery of bioorthogonal metabolic tracers. Looking forward, we anticipate several applications for the specific SCFA pro-metabolites demonstrated here. First, the strong labeling afforded by **6** suggest it may be a useful in vivo tracer for identifying endogenously short chain fatty acylated proteins associated with SCFA-driven phenotypes such as differentiation, anti-inflammation, or tumor suppression.^4, 6–9^ To be most impactful, such studies will require that pro-metabolites recapitulate the phenotypic effects of endogenous SCFAs. As many of these effects have been associated with the HDAC activity of SCFAs, it is promising that our cellular studies indicate the azido- and alkynyl- SCFAs investigated here also stimulate histone acetylation (Figure S2). Second, bioorthogonal pro-metabolites may provide a facile readout of global KAT/KDAC activity in cellular or in vivo systems, complementing current chemical proteomic approaches, which are only applicable in cell lysates.^46–47^ Finally, bioorthogonal SCFA pro-metabolites may facilitate chemical genetic approaches to identify specific KAT substrates in cells.^48^ Of note, mutations have been reported that confer KAT2A (GCN5L2) and KAT8 (MOZ) with the ability to use elongated azido- and alkynyl-CoAs.^49^ Overall, these studies represent a key methodological advance in the development of new methods to understand acetylation-dependent signaling in living systems.

## Methods

### General synthetic procedures and materials

Chemicals were purchased from commercial sources (Sigma-Aldrich, Alfa Aesar, and TCI America) and used without further purification unless otherwise noted. 2-azidoacetate (**1**), ethyl azidoacetate (**2**), 4-pentynoic acid (**9**), ethyl pentynoate (**10**), hexynoic acid (**13**), and ethyl hexynoate (**14**) were obtained commercially. 3-azidopropionic acid (**5**), carnitine esters (**3, 7, 11, 15**) and glycerol esters (**4, 8, 12, 16**) were prepared according to literature procedures.^50–52^ Flash column chromatography was performed using normal phase on a CombiFlash^®^ Rf 200i (Teledyne Isco Inc). ^1^H NMR spectra were recorded at 400 or 500 MHz, and are reported relative to deuterated solvent signals. Data for ^1^H NMR spectra are reported as follows: chemical shift (δ ppm), multiplicity, coupling constant (Hz), and integration. ^13^C NMR spectra were recorded at 100 or 125 MHz. Data for ^13^C NMR spectra are reported in terms of chemical shift. Analytical LC/MS was performed using a Shimadzu LCMS-2020 Single Quadrupole utilizing a Kinetex 2.6 μm C18 100 Å (2.1 × 50 mm) column obtained from Phenomenex Inc. Runs employed a gradient of 0→90% MeCN/0.1% aqueous formic acid over 4 minutes at a flow rate of 0.2 mL/min. Microwave experiments were carried out using the Biotage Initiator 4.0.1.

### General procedures and materials for cellular assays

A549 and HEK-293 cells were obtained from the NCI Tumor Cell Repository, while HepG2 cells were obtained from ATCC (Manassas VA). All cell lines were cultured at 37 °C under 5% CO2 atmosphere in a growth medium of RPMI supplemented with 10% FBS and 2 mM glutamine, with the exception of HEK293T cell lines, which were cultured in DMEM supplemented with 10% FBS, 2 mM glutamine. Cells were harvested by scraping and cell lysates prepared by sonication as previously described.^53^ Protein concentrations were determined by Qubit Protein Assay kit (Life Technologies #Q33212). Fluorescent labeling analyses were performed via Cu-catalyzed ligation to a TAMRA-azide or alkyne as previously described.^46^ SDS-PAGE was performed using Bis-Tris NuPAGE gels (4-12%, Invitrogen #NP0322), and MES running buffer (Life technologies #NP0002) in Xcell SureLock MiniCells (Invitrogen) according to the manufacturer’s instructions. SDS-PAGE fluorescence was visualized using an ImageQuant Las4010 Digital Imaging System (GE Healthcare). Total protein content on SDS-PAGE gels was visualized by Blue-silver Coomassie stain.^54^ SILEC experiments were performed as previously described. Acyl-CoAs were isolated by acid extraction and analyzed by LC-MS as previously described.^55–56^ 3-azidopropionyl-CoA was synthesized and purified by standard methods,^53, 57^ and used as a standard in LC-MS analyses. Histones were isolated by acid extraction and analyzed by immunoblotting as previously described.^33, 58^ Toxicity of pro-metabolites were assessed by sulforhodamine B staining^59^ after treatment for the dose/duration indicated. For proteomic analyses, HEK-293 proteomes (8 mg per replicate, treated with **6** or vehicle DMSO) were ligated to a biotin-alkyne using Cu-catalyzed [3+2] cycloaddition.^46, 60^ Enrichment, tryptic digest, LC-MS/MS analyses, and database searching was performed as previously described.^46^

### Treatment of cells with pro-metabolites for metabolic labeling analyses

For labeling experiments, cell lines were plated and allowed to adhere overnight. Cells were then treated with pro-metabolites **1-16** by adding of DMSO stock solutions directly to growth medium at the specified concentration, followed by gentle agitation and incubation for the specified time. For low glucose experiments, cells were grown in DMEM supplemented with 1 mM glucose (prepared from glucose-free media) for 24 h prior to addition of pro-metabolite **6**. For acetate and SAHA experiments, acetate (1 mM) or SAHA (10 μM) were added to media immediately prior to pro-metabolite **6**. In all experiments, final DMSO concentrations were < 0.5%. Unfractionated proteomes were harvested by washing adherent cells (80-90% confluent) 3x with ice cold PBS, and scraping cells into a Falcon tube followed by centrifugation (500 rcf × 5 min, 4 °C). Lysates were isolated from cell pellets by sonication as described above. Specific experimental treatment conditions are indicated in the figure captions.

### Mass spectrometry characterization of azidopropionyl-CoA formation

Cellular formation of azidopropionyl-CoA was confirmed by LC-HRMS and LC-MS/HRMS using previously described methods.^61^ Briefly, azidopropionyl-CoA (C_24_H_39_N_10_O_17_P_3_S), generated synthetically or by cell culture treatment in HepG2 cells using **5** or **6** was observed as an [M+H]^+^ (predicted *m/z* 865.1501, observed in cell culture extract 865.1505 dppm = 0.46) (Fig. S3b-c). Predominant fragmentation in MS/HRMS was derived from the neutral loss of 507 mass units from the loss of the adenosine with the pantetheine ejection maintaining the charge ([C_14_H_24_N_5_O_4_S]^+^ product predicted *m/z* 358.1544, observed 358.1543 dppm= −0.27) (Fig. S3a, Fig. S3c). To further confirm this finding, we performed stable isotope labeling by essential nutrients in cell culture (SILEC), treating Hepa1c1c7 murine hepatocellular carcinoma cells with ^13^C_3_ ^15^N_1_ pantothenate as previously reported.^62^ Treatment of cells with **6** (0.1 mM, 1 h) generated ^13^C_3_ ^15^N_1_-azidopropionyl-CoA, where the stable isotope labeling is enriched in the region of the CoA backbone generated from pantothenate. A mixture of 1:1 by volume of cell extract from unlabeled and SILEC labeled cells produced a perfectly co-eluting HRMS and MS/HRMS peak, validating our identification of azidopropionyl-CoA as a biochemically produced acyl-CoA metabolite (Fig. S3d).

## Acknowledgements

We thank the Laboratory of Proteomics and Analytical Technology for LC-MS/MS analyses and Dr. Hans Luecke (NIDDK) for helpful discussions. This work was supported by the Intramural Research Program of the NIH, National Cancer Institute (ZIA BC011488-02).

## Supplementary Material

**Figure S1.**
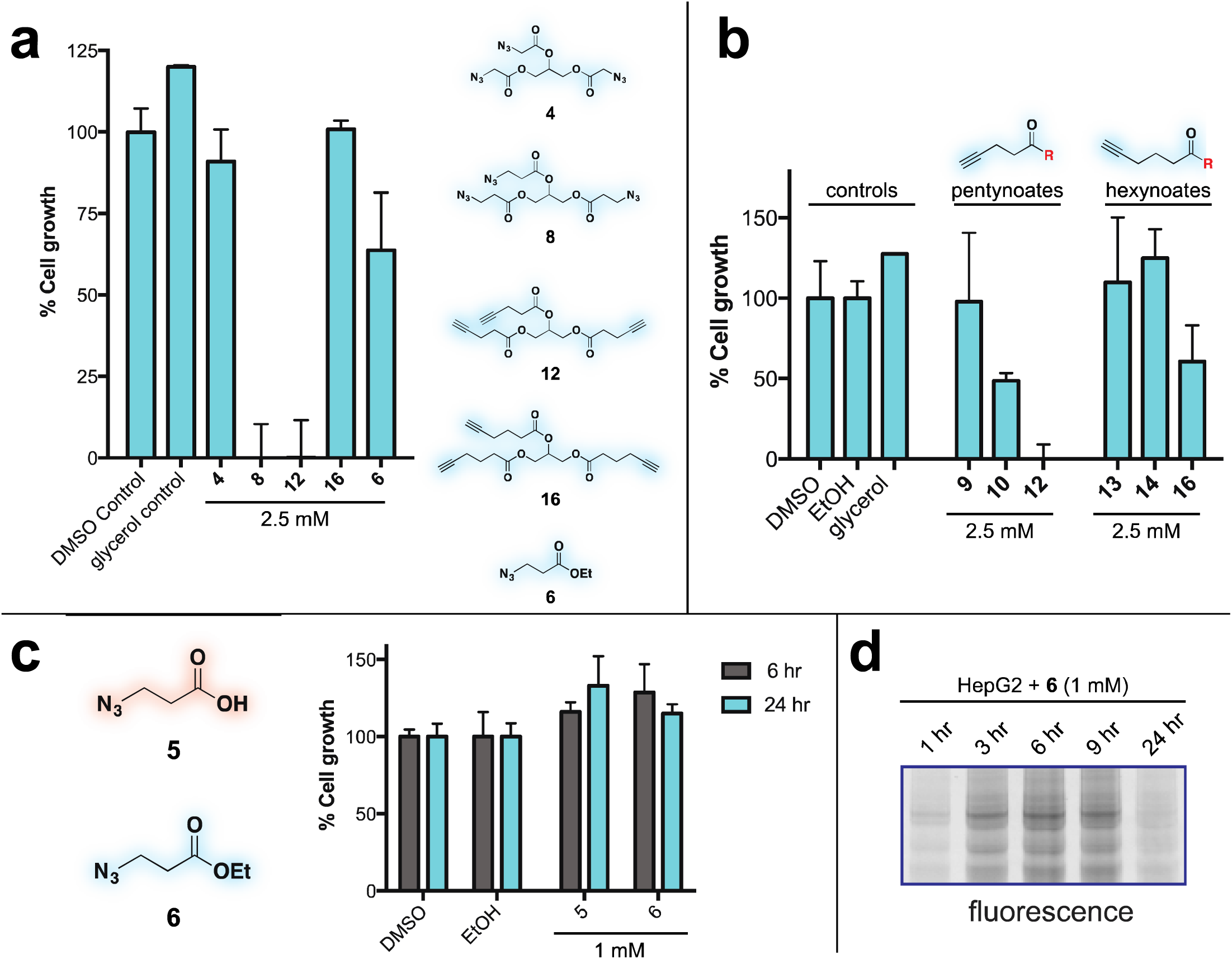
(a) Toxicity of active pro-metabolites at 24 h. (b) Relative toxicity of pentynoate and hexynoate analogues at 48 h. (c&d) Pro-metabolite **6** can be applied under conditions (1 mM, 6 h) that minimize cell death and enable maximal protein labeling.

**Figure S2.**
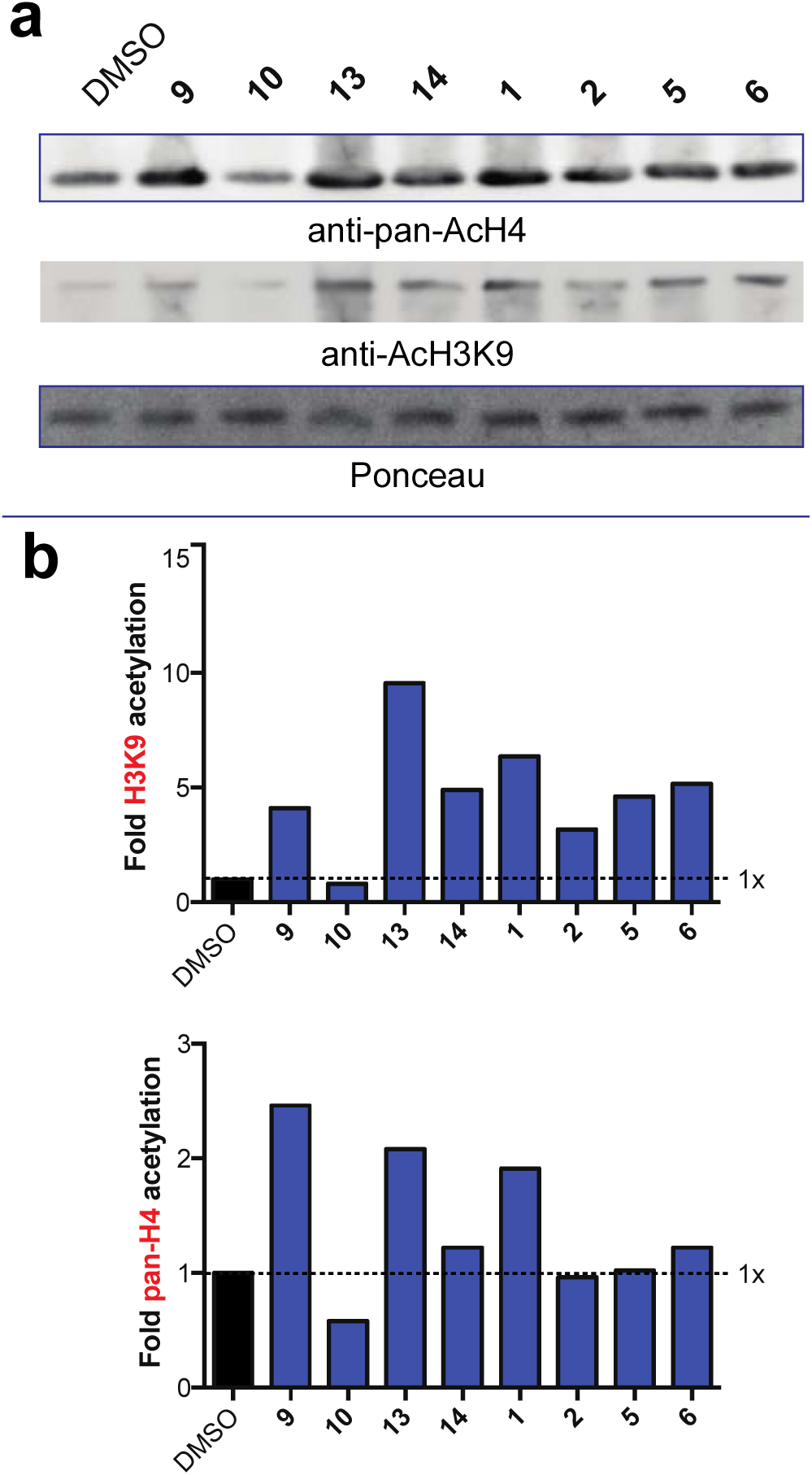
HDAC inhibitory activity of ethyl ester pro-metabolites and free acids. (a) Western blots of histone acetylation in HEK-293T cells following 6 h treatment with bioorthogonal acids (**1, 5, 9, 13**) or bioorthogonal esters (**2, 6, 10, 14**) at 2.5 mM. (b) Bar chart illustration depicting observed histone acetylation change.

**Figure S3.**
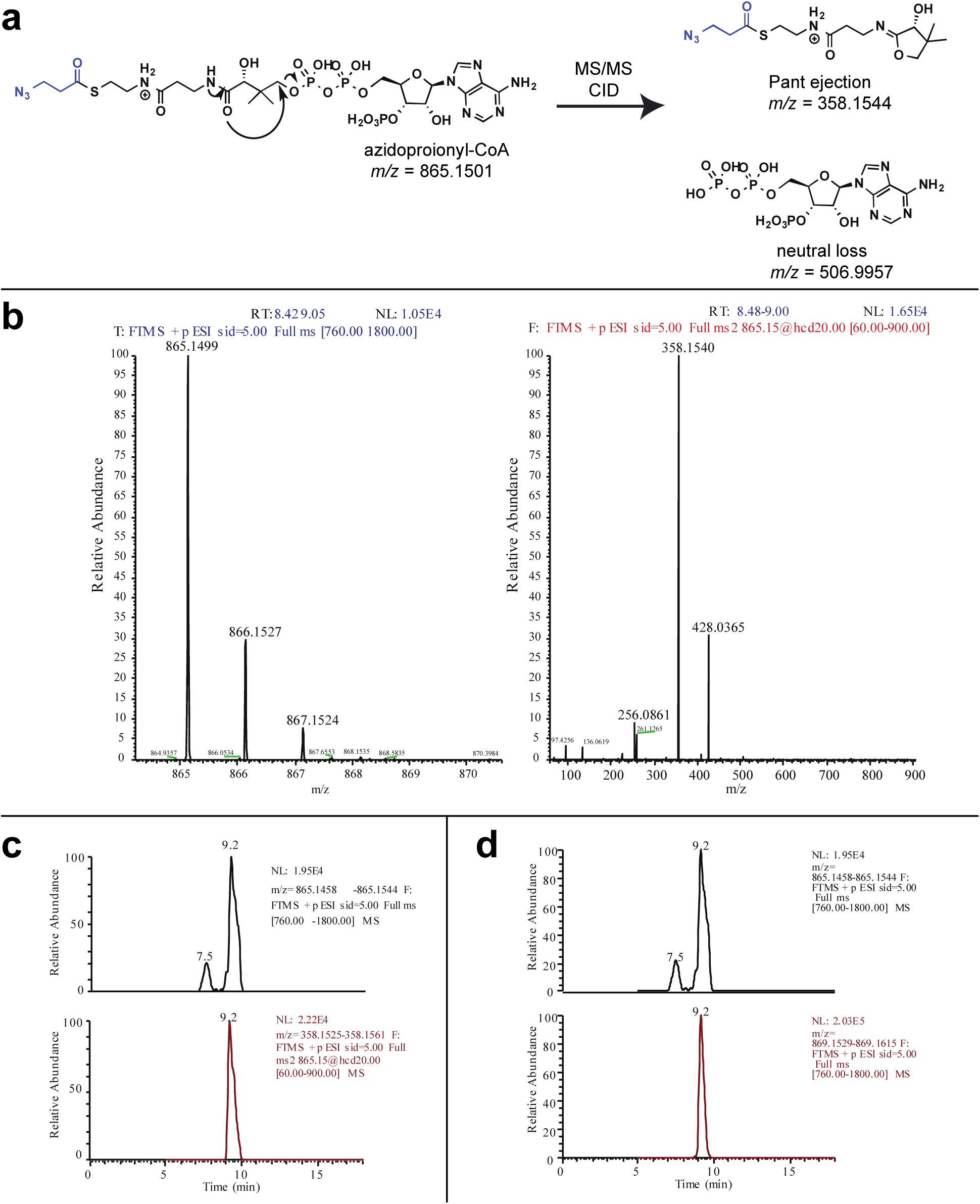
LC-MS/MS analysis of azidopropionyl-CoA formation. (a) Product ions derived from acyl-CoA analogues through collision-induced dissociation and MS/MS analysis. (b) HRMS (left) and MS/MS (right) of synthetically prepared azidopropionyl-CoA. (c) HRMS and LC-MS/MS of Hepa1c1c7 cell extract demonstrating co-eluting fragments specific to azidopropionyl-CoA. Top: 867.15 *m/z* corresponding to 3-azidopropionyl-CoA. Bottom: 358.15 *m/z* corresponding to pantetheine ejection fragment. Cells were treated with 0.1 mM ethyl azidopropionate (*6*) for 1 hr. Analogous results were observed in MCF7 and HepG2 cells treated with 6. (d) LC-HRMS demonstrating co-elution of azidopropionyl-CoA and ^13^C_3_ ^15^N_1_-azidopropionyl-CoA (9.2 min), which are well resolved from a near isobaric interference at 7.5 min.

